# Immediate TMS-EEG responses reveal motor cortex excitability

**DOI:** 10.1101/2024.08.20.608770

**Authors:** Antonietta Stango, Agnese Zazio, Guido Barchiesi, Natale Salvatore Bonfiglio, Elisa Dognini, Eleonora Marcantoni, Marta Bortoletto

**Author notes:** Equal contribution (Antonietta Stango, Agnese Zazio).

## Abstract

**Background:** Combined transcranial magnetic stimulation and electroencephalography (TMS-EEG) is widely used to probe cortical excitability at the network level, but technical challenges have prevented its application to investigate local excitability of the stimulated area. A recent study revealed immediate TMS-evoked potentials (i-TEPs) after primary motor cortex (M1) stimulation, suggesting that it may represent a local response. Here, we aimed at testing if this activity is physiological in nature and what it represents.

**Methods:** We analyzed a TMS-EEG dataset from 28 healthy participants recorded at 9.6 kHz including two M1 stimulation conditions with opposite biphasic current directions. We localized the brain sources of i-TEPs, calculated the immediate TMS-related power (i-TRP) to distinguish between two oscillatory components that may contribute to i-TEPs, and investigated the relationship between i-TRP and motor-evoked potentials (MEPs). In an additional recording, we stimulated a control site evoking a muscular response to understand the contribution of the TMS-related muscle artifact.

**Results:** Results confirmed i-TEPs with similar characteristics as previously described. The i-TRP revealed strong activity in two ranges 600-800 Hz and 100-200 Hz; The former was positively associated with MEPs amplitude for both current direction conditions. Moreover, i-TEPs were localized in the precentral gyrus of the stimulated hemisphere and the muscular response generated by the control stimulation site differed from i-TEPs and i-TRP.

**Discussion:** These findings provide first evidence on the physiological nature of i-TEPs and i-TRP following M1 stimulation and that i-TRP represents a direct measure of excitability of the stimulated cortex.

## Introduction

Concurrent transcranial magnetic stimulation-electroencephalography (TMS-EEG) has been proposed as a non-invasive painless technique to study excitability of cortical networks in healthy populations as well as in clinical conditions, such as neuropsychiatric disorders (Tremblay et al. 2019; Farzan 2024). Accordingly, current TMS-EEG measures of cortical excitability, including both TMS-evoked potentials (TEPs) (Ilmoniemi et al. 1997) and event-related spectral perturbation (Rosanova et al. 2009), reflect activations in cortical networks including both the target and connected areas (Massimini et al. 2005; Bortoletto et al. 2015; Ozdemir et al. 2020; Momi et al. 2021).

A local excitability index of the stimulated area would allow a non-invasive assessment of the cortical inhibition/excitation balance, advancing current understanding and treatment of various neuropsychiatric conditions, but it has not been identified yet due to methodological challenges. In fact, the EEG signal in the first milliseconds after the TMS pulse, when a local immediate response most likely occurs, is covered by several artifacts, as the TMS artifact, the decay artifact and the muscle artifact (Hernandez-Pavon et al. 2023b). Therefore, the first 5-10 ms after TMS (Veniero et al. 2009) are removed during offline signal processing. When the signal is recovered, the responses already include secondary activations of the areas surrounding the target and of more distant areas (Ilmoniemi et al. 1997; Bortoletto et al. 2021; Zazio et al. 2021, 2022; Guidali et al. 2023; Hernandez-Pavon et al. 2023a).

In a recent paper (Beck et al. 2024), EEG signal was recovered at approximately 2 ms after the TMS pulse and showed new TEP components occurring before 6 ms. The authors highlighted that these immediate components (i-TEPs) included high-frequency activity around 600-800 Hz, resembling the corticospinal descending volleys recorded in the spinal cord, and suggested that they may represent a local response of the stimulated cortex. To support this claim, they excluded that i-TEPs may be an electromagnetic artifact as they were not observed for stimulation with the same parameters over two parietal control areas. Moreover, they reported that the amplitude of i-TEPs was modulated by current direction as well as by the intensity of TMS. However, the effect of current direction on I-waves for biphasic stimulation has not been directly tested (Di Lazzaro et al. 2001; Ziemann 2020), and the effect of intensity of stimulation, although reported for I-waves (Burke et al. 1993; Kaneko et al. 1996; Di Lazzaro et al. 1999), does not fully rule out the possibility that i-TEPs are an artifactual activity. For example, the muscle artifact would also increase with stimulation intensity. Eventually, there is no indication that the activity occurring in the i-TEP window represent motor excitability.

Here, the goal of the present work was to understand the origin of the immediate TMS-EEG responses. First, we aimed to confirm the recent finding on i-TEPs, i.e., to record the i-TEPs for AP-PA and for PA-AP current direction, with a different TMS-EEG system; Second, we aimed to clarify whether immediate TMS-EEG responses have a cortical or an artifactual origin, controlling for the contribution of muscle artifacts. Finally, we calculated the immediate TMS-related power (i-TRP) in high-frequency bands as the event-related spectral perturbation that includes both evoked and induced activity, and we tested its relationship with a typical peripheral measure of cortico-spinal excitability, i.e., motor-evoked potentials (MEPs).

## Methods

### Main experiment

#### Participants

The data presented in this paper were collected at the Neurophysiology Laboratory of the IRCCS Istituto Centro San Giovanni di Dio Fatebenefratelli (local ethics committee reference number: 102-2021) and are part of the dataset published in the work by Guidali et al. (2023) to measure the modulation of a positive component occurring in the contralateral hemisphere at about 15 ms after the TMS and it can be found at https://gin.g-node.org/Giacomo_Guidali/Guidali_et_al_2023_EJN_RR. In the original study, 28 right-handed participants from 18 to 50 years old completed the study with stimulation intensity below 90% of the maximal stimulator output (MSO) in all conditions.

#### Data acquisition

TMS-EEG was acquired from 74 EEG electrodes at 9.6 kHz using the g.HIamp amplifier (g.tec medical engineering GmbH). This sample rate is sufficient with this EEG system to record a TMS artifact of about 2 ms, as shown by data from our lab (Zazio et al. 2022; Guidali et al. 2023)and in Freche et al. (2018). The ground electrode was placed on the tip of the nose and reference was FPz. Skin-electrode impedance was below 5 kΩ. Before and during recording, the signal was visually checked as continuous EEG and averaged TEPs to monitor for artifacts, such as prolonged decays, noisy channels, and line noise. EMG was recorded at the same sampling rate from the right *abductor pollicis brevis* using a bipolar belly tendon montage.

Single pulse TMS was delivered with a figure-of-eight coil (Magstim model Alpha B.I. Coil Range, diameter: 70 mm) over the left primary motor cortex (M1) at 110% of the resting motor threshold (rMT). The original study included three current directions for both monophasic and biphasic pulse waveforms, but here only biphasic stimulation (Magstim Rapid^2^, Magstim, Whitland, UK) was considered. Specifically, the 2 current directions previously reported by Beck et al were considered: AP-PA, PA-AP. Coil position was monitored using SofTaxic Optic 3.4 neuronavigation software (EMS, Bologna, Italy; www.softaxic.com) throughout recordings. Table 1 reports rMT for each subject and for each condition. Further information can be found in Guidali et al. (2023).

**Table 1.**
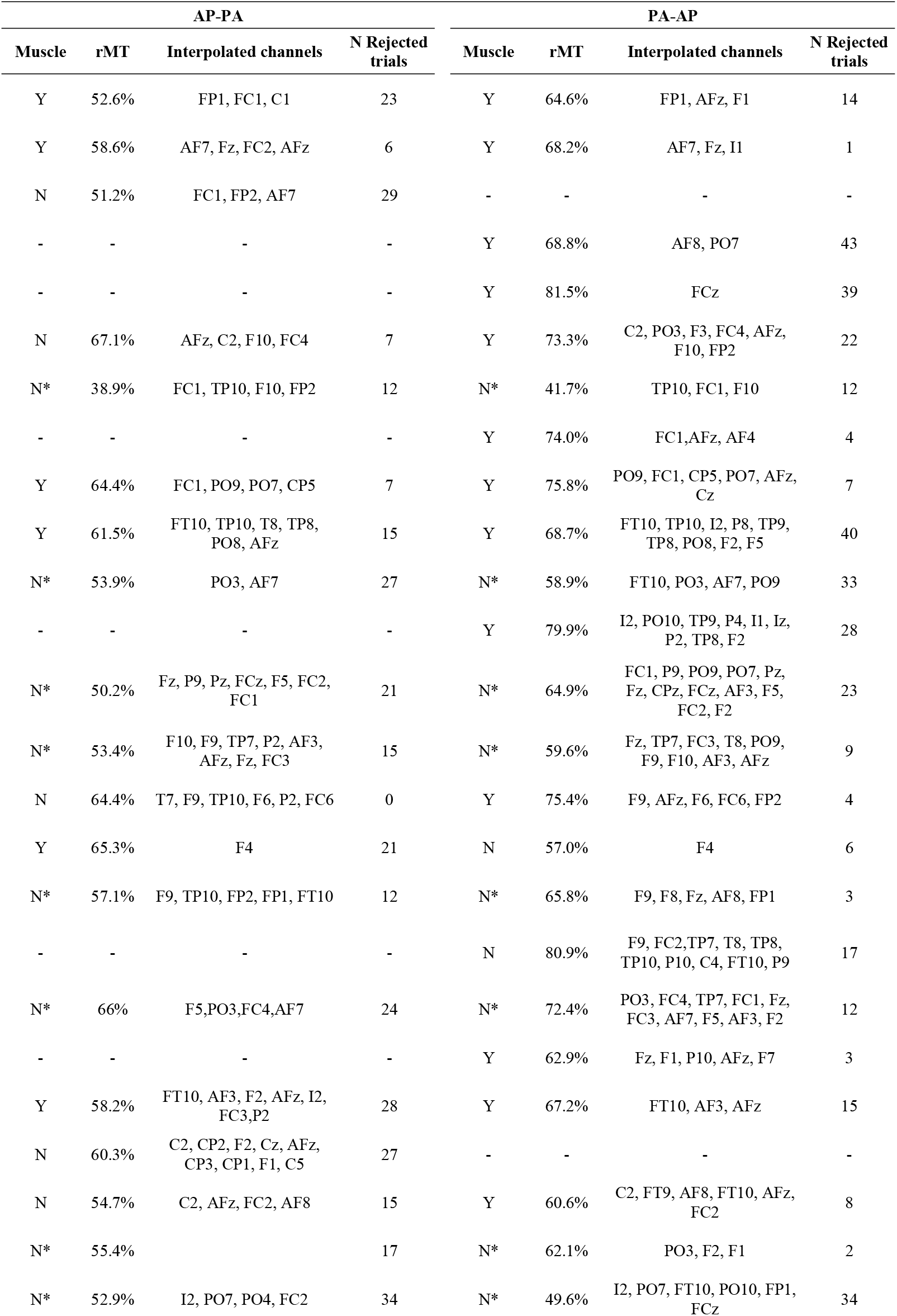
Individual data on rMT, presence of muscle artifact, interpolated channels and number of rejected epochs for both AP-PA and PA-AP conditions. Missing data refers to cases which were excluded from further analyses because of decay artifacts or noise. Asterisks indicate subjects that were included in the final sample for statistics, as they belonged to the noMuscle groups for both AP-PA and PA-AP conditions.

#### Preprocessing of i-TEPs, i-TRP and MEPs

To obtain i-TEPs, the preprocessing was limited to epoching with baseline correction, artifact rejection, and identification and interpolation of channels with strong decay. The signal between 0 and 2.8 ms was removed. The artifact rejection was manually run to remove ocular artifacts and noisy trials, and was applied on epochs from -500 ms to 500 ms. At this stage, electrodes with decay artifacts were identified and interpolated (Table 1). Then, we obtained shorter epochs, from -100 ms to 100 ms, and applied a baseline correction in the first 95 ms, i.e. from - 100 ms to -5 ms.

Subjects were excluded from further analyses if EEG signal was contaminated by decay artifacts or was noisy in the window 2-6 ms. After this selection, 21 subjects were kept for AP-PA condition, 25 for PA-AP condition. Finally, data were averaged to obtain i-TEPs and, for each current direction, subjects were divided in two groups, i.e., NoMuscle and Muscle, based on the presence of a visible muscle artifact in the averaged signal. Muscle artifact was identified by visual inspection as a MEP-like response with a lateralized topography in the stimulated hemisphere. The NoMuscle groups included 14 subjects in the AP-PA condition and 10 subjects in the PA-AP condition. The Muscle group included 7 subjects in the AP-PA condition and 15 subjects in the PA-AP condition. A total of 8 subjects belonged to the noMuscle group for both AP-PA and PA-AP conditions.

For the i-TRP, we run time-frequency representation analysis and computed power on single-trial data from 30 ms before to 100 ms after the TMS pulse in a frequency range from 30 to 1000 Hz. Hanning taper was applied on a frequency-dependent time window of five cycles per frequency, sliding in steps of 1 ms. Output measures were restricted to C3 as the closest channel to M1 hotspot. TRP was averaged for each trial in the two time-frequency windows presenting maximal activity: Between 3 and 6 ms as average of 600 and 800 Hz (fast, fi-TRP), and at the same time-window as average of frequencies between 100 and 200 Hz (slow, si-TRP).

To measure MEP amplitude, continuous EMG was cut around the TMS pulse, i.e., from 0 ms to 3 ms (TESA cubic interpolation), bandpass-filtered between 10 and 2500 Hz and notch-filtered at 50 Hz (FIR filtering) and epoched from 200 ms before to 500 ms after the TMS pulse. Epochs with muscular or background noise, as indicated by amplitude exceeding 100 μV in the 50 ms before the TMS pulse, were rejected. Finally, MEP amplitude in each trial was calculated as peak-to-peak difference in the interval between 20 and 50 ms after TMS.

At the end of preprocessing, only trials with both TEPs and MEPs were included in the final dataset: the number of trials considered for each subject ranged between 46 and 68 for AP-PA condition and between 46 and 78 for PA-AP condition.

Preprocessing was performed in Matlab (The Mathworks, Natick, MA, USA) using EEGLAB functions (Delorme and Makeig 2004) and Fieldtrip (Oostenveld et al. 2011).

#### Source localization

We used BESA Research (v. 7.1, BESA GmbH, Gräfelfing, Germany) to create a source model of the i-TEPs based on dipole fitting. The source model was computed separately for the AP-PA condition and for the PA-AP condition. The procedure started by fitting one dipole to the grand-average i-TEPs of the NoMuscle group in the interval between 2.8 and 6 ms. In this way, we optimized the signal-to-noise ratio during the dipole fitting by reducing the presence of artifacts superposed to the signal of interest. The percentage of the residual variance (RV) in the time-window of interest, i.e., 3-6 ms, was considered as an estimate of the goodness of fit of the model and to define if more dipoles had to be added. Once the dipole model was completed for the NoMuscle group, it was used as the starting point to create the dipole model in the Muscle group. Again, the RV was considered to decide whether the model sufficiently explained the data or if it needed to include more dipoles. In the latter case, dipoles were fitted at the time of the peak of residual principal components. Dipoles were fitted until RV reached values below 20% in the window 2.8-6 ms. Finally, we obtained the position of the dipoles as MNI coordinates, corresponding nearest anatomical structure and associated Brodmann area.

#### Statistics

Statistics were run only in the sub-group of subjects that had been included in the NoMuscle group for both AP-PA and PA-AP conditions. In this way, effects could be investigated in a within-subject design. Generalized mixed models with a gamma distribution and a log link function were used to test the relationship between single-trial i-TRP (as a fixed effect) and MEP amplitude (as the dependent variable), including the interaction between i-TRP and current direction. Subject was included as a random effect, and current direction (AP-PA, PA-AP) was included as a fixed effect. Separate models were run for fi-TRP and si-TRP. Moreover, generalized mixed models (gamma distribution, log link function) with subject as random effect were used to test for changes in i-TRP (fi-TRP and si-TRP as dependent variables in separate models) depending on current direction (AP-PA, PA-AP; as fixed effect). Statistics were run in R v. 4.3.2 (R Core Team, 2021) using the packages ‘lme4’ for mixed-effects models (Bates et al. 2015) and ‘emmeans’ for post-hoc comparisons (Lenth et al. 2024).

### Control experiment

To further exclude the contribution of the muscle artifact to the i-TEPs, we ran a control experiment in which aimed at evoking cranial muscles’ responses without cortical stimulation, by delivering TMS pulses over a lateral site in correspondence with the temporal branch of the facial nerve at a very low intensity. Then, we compared the obtained signal with i-TEPs obtained from the stimulation of the cortical motor hotspot (Figure 3A).

TMS-EEG was acquired in one subject from C3 at 9.6 kHz (g.HIamp amplifier g.tec medical engineering GmbH). The ground electrode was placed on the tip of the nose and reference was FPz. Skin-electrode impedance was below 5 kΩ. The signal was visually checked during recording to monitor for artifacts. EMG was recorded at the same sampling rate from the right *first dorsal interosseous* using a bipolar belly tendon montage. For each condition, 100 single TMS pulses were delivered at random intervals with a figure-of-eight coil (Magstim model Alpha B.I. Coil Range, diameter: 70 mm). Coil position was monitored using SofTaxic Optic 3.4 neuronavigation software (EMS, Bologna, Italy; www.softaxic.com) throughout recordings.

To obtain i-TEPs, TMS was delivered over the left primary motor cortex (M1) at 110% of the resting motor threshold (rMT), i.e., 59% of the MSO, with 45 degrees coil orientation to the medial-sagittal plane and AP-PA current direction.

To evoke cranial muscle twitches while minimizing cortical stimulation, we aimed at cranial nerves with low intensity stimulation. This procedure was based on evidence that nerves are more excitable than muscles and their stimulation can induce muscle contraction within a few ms (Purves-Stewart and Worster-Drought 1952). Therefore, TMS intensity was set at 10% of the MSO and the coil was moved laterally to C3, on the left side of the scalp. The nerve hotspot was defined as the location that evoked the biggest muscle artifact in C3. Once the hotspot was localized, the coil position was kept constant and TMS intensity was varied. We started from 5% of MSO and increased intensity at 10%, 13% and 15% of MSO so that a clear muscle artifact could be seen in C3.

Data were preprocessed to obtain i-TEPs and i-TRP over C3 with the same procedure of the main experiment, except that data were epoched from -50 to 50 ms.

## Results

### i-TEPs

In both current direction conditions, a series of three peaks superposed to a positive slower potential was visible within the first 6 ms after TMS. The topographic distribution revealed higher amplitude around the C3 electrode. A slightly delayed latency compared to previous finding (Beck et al. 2024) is explained by a different interval between TMS marker and TMS artifact, so that in our system the TMS marker starts 1 ms before the TMS pulse (Figure 1A-B). These peaks were visible also in the averaged signal of single subjects, although with some variability (Figure S1-S2), and even at single-trial level in the NoMuscle group (Figure S3-S4). Therefore, the recorded signal resembled the i-TEPs reported by Beck et al. 2024.

**Figure 1.**
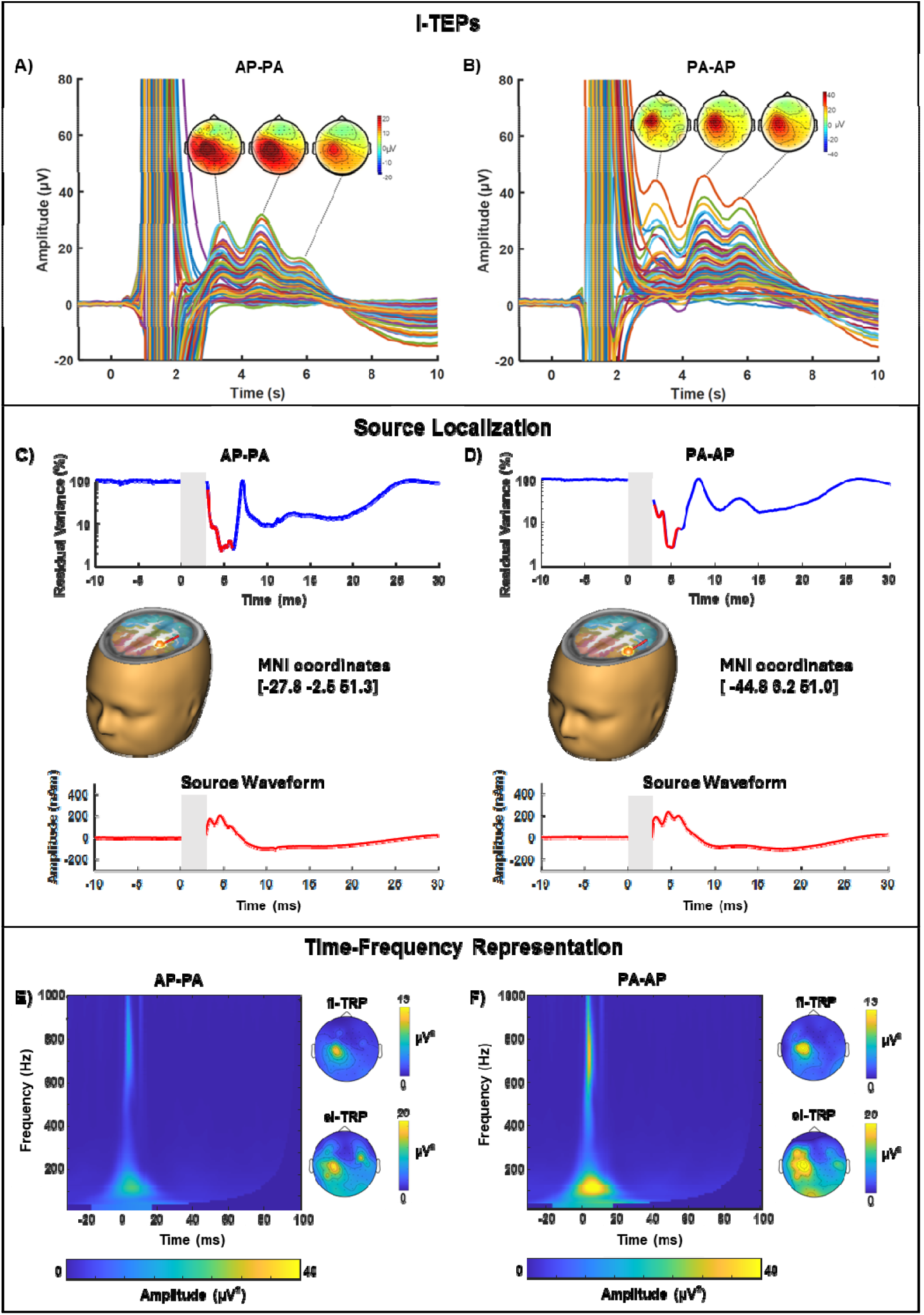
Grand-average across trials of the NoMuscle groups for AP-PA (panels A, C, E) and PA-AP (B, C, D) conditions. The top panels (A, B) depict the i-TEPs and the topography at the three peaks; the TMS artifact was interpolated in the analyses and is shown here for display purposes. A short delay (< 1 ms) of the TMS pulse with respect to the TMS marker (i.e., 0 ms on x axis) explains the slight difference in pea latencies compared to Beck et al. (2024). The middle panels (C, D) show the source localization of the dipole that best explains the signal recorded over the scalp. The bottom panels (E,F) show the i-TRP at channel C3 and the topographies at the two time-frequency windows of interest (amplitude range in colorbars).

To understand the contribution of the TMS-induced muscle artifact, we also explored data of subjects where this artifact was visually detected (i.e., Muscle group). In the AP-PA condition, the muscle artifact had generally low amplitude and the characteristic three peaks of i-TEPs were still visible in the grand-average (Figure S5A-B) and in single subject data of many subjects (Figure S6-S7). In the PA-AP condition, the muscle artifact had a stronger impact on the signal, possibly due to the higher stimulation intensity employed in this condition.

### Source localization

The time-varying cortical activity that explains the scalp i-TEPs is shown in Figure 1C-D. As can be seen, grand-average i-TEPs for the NoMuscle groups were substantially explained by a single dipole (Dip1) localized in the left precentral gyrus for both current directions, corresponding to Brodmann area 6 (MNI coordinates, AP-PA: -27.8, -2.5, 51.3; PA-AP: -44.8, 6.2, 51.0). The RV of these models for the interval of interest was 8% for AP-PA and 9% for PA-AP, although higher RV values are present soon after the TMS pulse, possibly due to residual decay artifacts. Interestingly, the temporal pattern of the dipole activity showed the three peaks characteristic of i-TEPs.

For Muscle groups, the source analysis resulted in two different models depending on current direction. In the AP-PA condition (Figure S5), adding one more dipole (Dip2) to the source model was sufficient to obtain a RV of 13%, over the interval of interest. Dip2 was located outside the brain, on the lateral aspect of the head, just in front of the left ear (MNI coordinates: -67.5, 16.8, -42.7). The dipole’s placement was in line with muscles located on the lateral side of the head, as auricularis muscles or the temporalis muscle. Importantly, the time course of activity of Dip2 resembled a M-wave (Di Bella et al. 1997), while the three peaks typical of i-TEPs were still visible in Dip1.

In the PA-AP condition (Figure S5), the muscle artifact was more prominent and the model required 2 more dipoles (Dip2 and Dip3) to reach a RV below 20%, i.e. 9%. Dip2 was positioned again on the left side of the head, in front of the ear (MNI coordinates: -80.2, 7.9, -20.1). Dip3 was positioned in between Dip1 and Dip2, outside the edge of the cortical surface (MNI coordinates: -68.3, 6.46, 22.9). The activity modeled by these two dipoles was strong and similar to MEPs, while Dip1 seemed to involve smaller activity and faster oscillations.

### i-TRP

The time-frequency representation analysis in the NoMuscle group showed increased power after TMS in two frequency ranges for both AP-PA and PA-AP conditions: Between 100-200 Hz and between 600-800 Hz (Figure 1E-F).

In the Muscle group (Figure S5E-F), the i-TRP profile was similar to the NoMuscle group for the AP-PA condition. However, in the PA-AP condition, in which the muscle artifact is stronger than cortical activity, the power was stronger, mainly in the lower frequencies.

### Relationship with MEPs and current direction modulation

The generalized mixed model showed that fi-TRP positively predicted MEP amplitude (*t*=3.2, *p=*0.001) so that the stronger the power in fi-TRP, the greater the MEPs (Figure 2A). Moreover, we found a significant effect of current direction indicating that MEPs were higher in AP-PA condition than in PA-AP (*t*=2.7, *p*=0.007). No significant interaction between fi-TRP and current direction was observed (*t*=-0.8, *p*=0.43).

**Figure 2.**
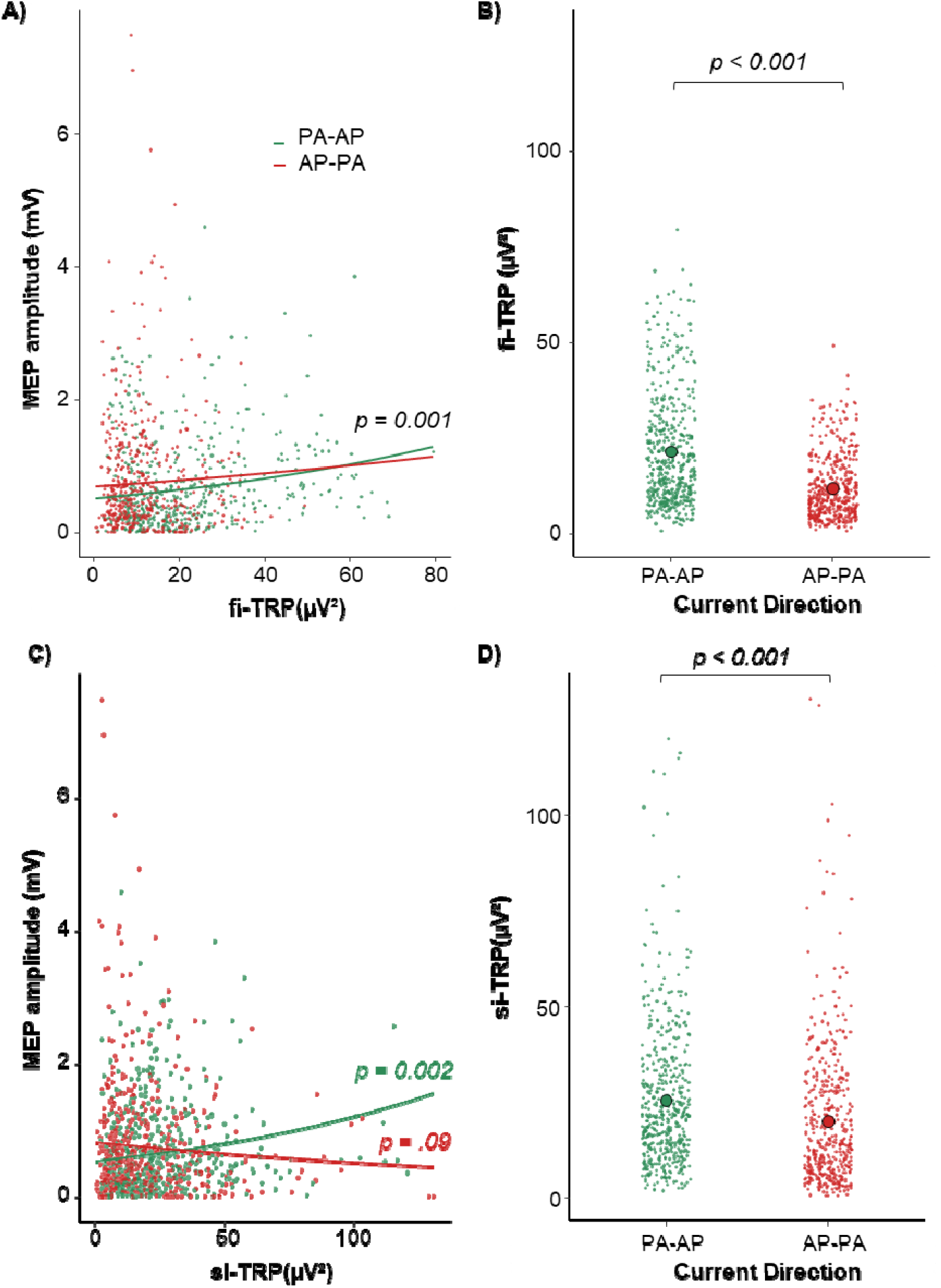
Relationship with MEPs and current direction modulation for fi-TRP (A and B, respectively) and si-TRP (C and D, respectively). Dots represent single trial values. Fitted lines in A represent the main effect of fi-TRP along with the absence of interaction between fi-TRP and Current Direction, while those in C reflect the interaction effect between si-TRP and Current Direction, with only PA-AP showing a significant positive relationship with MEP amplitude. Big dots in B and D indicate the average of i-TRP over trials.

**Figure 3.**
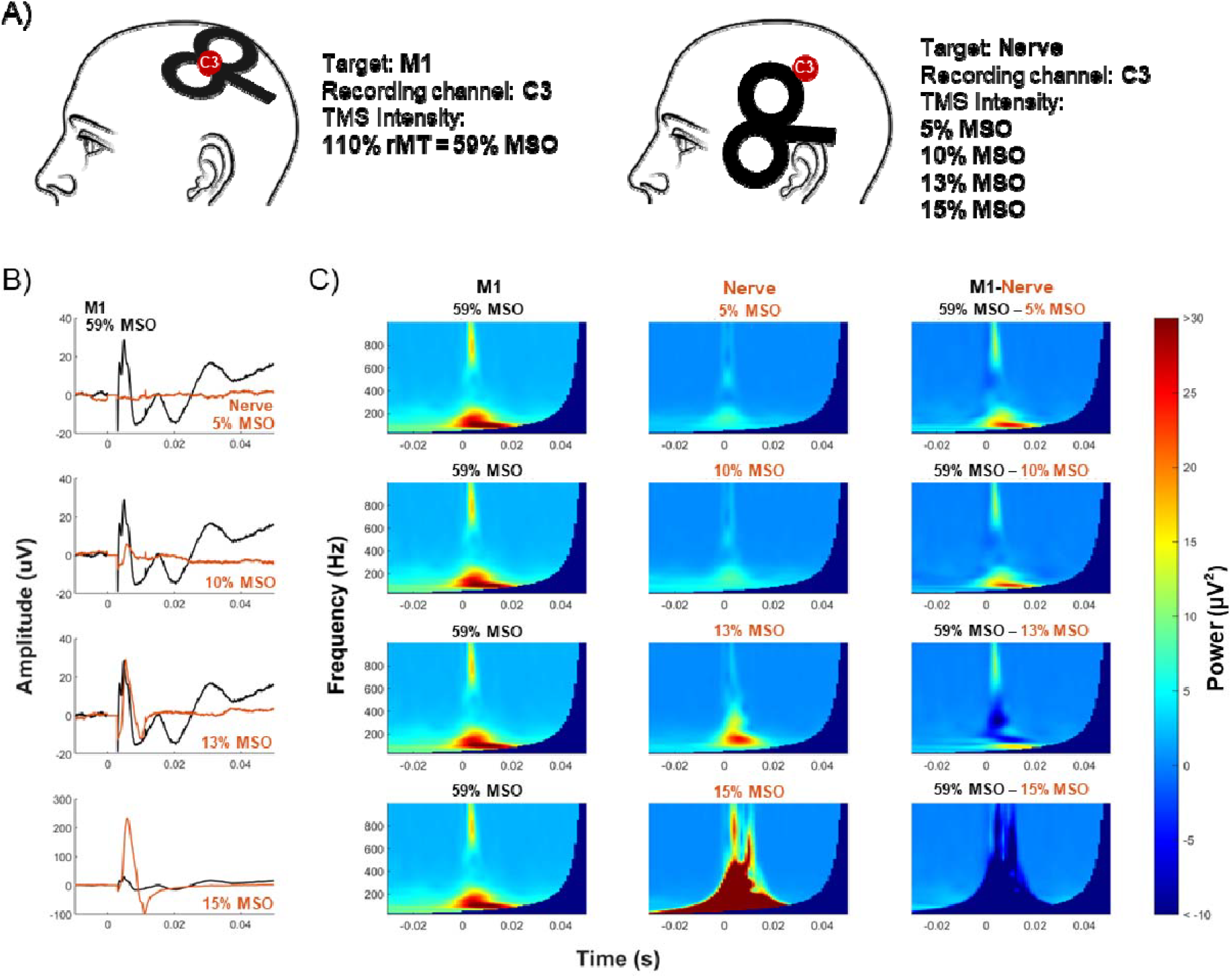
Control experiment. A) Schematic representation of the two stimulation conditions, targeting M1 hotspot (left) and the cranial nerve (right). B) Average TEPs recorded after M1 stimulation at 110% of rMT (black trace) overlapped to nerve stimulation (orange trace) with increasing intensity from top to bottom row (i.e., 5%, 10%, 13% and 15% of MSO). C) TFR of TEPs recorded in C3 after M1 stimulation at 110% of rMT (left column), after nerve stimulation with increasing intensity from top to bottom row (middle column) and the difference between the two (right column). Time frequency representation range in colorbar.

The same model including si-TRP as predictor also showed a positive relationship with MEPs (*t=*3.1, *p=*0.002), likely explained by PA-AP, as revealed by the significant si-TRP and current direction interaction (*t=*-3.6, *p<*0.001). Indeed, post-hoc comparisons showed a significant positive relationship between si-TRP power and MEPs amplitude in PA-AP (*p*=.002) but not AP-PA (*p=*0.09; Figure 2C). The effect of current direction on MEPs was confirmed (*t*=4.3, *p<*0.001).

Finally, we found that both fi-TRP (*t*=-19.8, *p*<0.001) and si-TRP (*t=-*7.5, *p*<0.001) were modulated by current direction and were higher in PA-AP compared to AP-PA (Figure 2B-D).

### Control experiment

The stimulation of the motor hotspot at 110% of the rMT resulted in visible i-TEPs in C3. Amplitude and latency were in line with data reported in the main experiment.

The stimulation of the nerve hotspot was located over FT7. Stimulations at 10% of MSO and above evoked a visible muscle twitch that increased with stimulation intensity (Figure 3B). Although the nerve stimulation at 13% of MSO evoked a muscle twitch of similar amplitude of i-TEPs, the former did not present the high-frequency peaks typical of i-TEPs. The time-frequency representation analysis showed visual differences for M1 stimulation and for nerve stimulation: the motor hotspot stimulation presented power increase in the two frequency ranges reported in the main experiment, i.e., 600-800 Hz and 100-200 Hz (Figure 3C). The nerve stimulation was associated with power increase in the range 100-200 Hz at lower intensity, and this range increased toward higher frequencies as stimulation intensity increased. Importantly, nerve stimulation at 13% of MSO, in which the muscle twitch had amplitude similar to i-TEPs, power was increased in frequencies ranging from 100 Hz to 400 Hz. The difference in the i-TRP revealed that M1 stimulation induced higher power than nerve stimulation in the range 600-800 Hz; Differently, nerve stimulation induced higher power than M1 stimulation at lower frequencies up to 500 Hz.

## Discussion

Our study confirms the presence of i-TEPs within the first 6 milliseconds post-stimulation, characterized by three peaks with a topographical distribution over centro-lateral electrodes surrounding M1 hotspot, for both AP-PA and PA-AP current directions. Most importantly, our data provide first evidence that activity recorded in the i-TEPs window is physiological in nature and represents the neuronal excitability of the stimulated motor cortex. First, we found a positive association between the i-TRP and the amplitude of MEPs. Second, source localization models supported that i-TEPs are generated in the precentral gyrus of the stimulated hemisphere and their activity can be clearly distinguished from sources of muscle twitches. Finally, the control experiment suggested that immediate TMS-EEG responses do not reflect muscle artifacts, as evidenced by the distinct pattern compared to muscle twitches.

In understanding the physiological origin of the activity recorded immediately after the TMS, we considered both phase-locked activity, i.e., the i-TEPs, and the total oscillatory response, i.e., the i-TRP, which comprises both phase-locked (i.e., evoked) and non-phase locked (i.e., induced) activity (Kalcher and Pfurtscheller 1995; Herrmann et al. 2014). Although TEPs and TRP do not necessarily represent the same activity, they are likely closely related in our study. Indeed, i-TEPs are visible as strong phase-locked activity at single trial level in all subjects, suggesting that phase-locked activity is stronger than induced oscillations and that i-TRP mainly represents i-TEPs. This is in line with previous evidence that TMS induces a short-lasting phase reset of ongoing oscillations (Romei et al. 2010, 2011; Herring et al. 2015).

Importantly, time-frequency analysis allowed us to disentangle the contribution of faster and slower components of i-TEPs both in the relationship with MEPs amplitude as well as in the comparison between current directions. Indeed i-TEPs appear to be composed by a fast oscillation superposed to a slower response, as suggested upon visual inspection by Beck et al. (2024). This intuition was confirmed in the time-frequency analysis that showed two clusters of increased power after TMS between 600-800 Hz and between 100-200 Hz in the first ms after the TMS pulse, on the electrode closest to stimulation site. The fast frequencies align with the estimated relevant frequencies for I-waves (Di Lazzaro et al. 2012) and may be generated by fast pyramidal tract neurons in the sensorimotor cortex, as they have been shown to respond to current injection with double or triple spikes and an interspike interval of about 1.5 - 2.5 ms (Calvin and Sypert 1976; Curio 2000). Despite EEG is more sensitive to postsynaptic potentials (Da Silva in Mulert and Lemieux 2023), the high synchronization induced by TMS at early latencies might enable the detection of action potentials using scalp electrodes (Di Lazzaro et al. 1998). Indeed, evidence of EEG sensitivity to synchronized action potentials has already been reported for EEG activity at ∼600 Hz during median nerve stimulation (Baker et al. 2003). Overall, these data support that the si-TRP represent the cortical precursor of I-waves.

The si-TRP, i.e., 100-200 Hz, falls within the “ripples” range. These oscillations have been observed in healthy brains in several brain regions, including sensorimotor areas of the neocortex, and in several cognitive states (Matsumoto et al. 2013; Gerner et al. 2020). Thus, the TMS coils act as an external stimulus, inducing activity that is physiologically plausible in the target area. While it is difficult at this point to speculate about the physiological mechanisms underlying the si-TRP, we hypothesize that it might reflect a hypersynchronization of intracortical postsynaptic potentials, that, being slower compared to action potentials, might maintain a synchronization for a longer period, but resolving within ∼20 ms.

The fast and the slow i-TRP showed a different pattern of relationship with MEPs: fi-TRP was positively associated with MEPs independently of current direction, whereas the association between the si-TRP and MEPs was significant for the PA-AP condition only. It should be considered that if the activity recorded in the cortex is a precursor of I-waves in the corticospinal tract and then MEPs in the muscles, it should be: a) present for both current directions; b) associated with MEPs for both current directions. Although the former is true for both si- and fi-TRP, the latter was verified for fi-TRP only.

The results on si-TRP may be explained by several factors. Considering that ripple generation is influenced by larger networks than very fast ripples, it is possible that a bigger network is more affected by changes in current directions. Alternatively, this frequency range may be more prone to the influence of concomitant muscle artifacts (Barchiesi et al. 2020). However, more studies are needed to understand the functional meaning of the si-TRP that we found in this study.

Furthermore, both fi- and si-TRP were modulated by current direction, such that PA-AP was stronger than AP-PA. Such finding is in line with what reported by Beck and colleagues on the amplitude of i-TEPs peaks 2 and 3 when adjusting the TMS intensity on the rMT of the respective current directions. On the other hand, MEPs amplitude was higher for AP-PA compared to PA-AP. This result is consistent with the hypothesis that different neural populations are activated by the two current directions, with AP-PA being the optimal one to activate the cortico-spinal tract (Siebner et al. 2022).

Another evidence that the immediate TMS-related activity has a cortical origin and is evoked in the area around the site of stimulation comes from the source localization of i-TEPs within the precentral gyrus. The dipole explaining cortical activation in cases with no muscular response maintained the characteristic three peaks unfolding in the first 3 ms of recovered signal, and this feature was also clearly visible in the AP-PA condition in which the muscle artifact was of relatively low amplitude. The location of the dipole resulted slightly more anterior and deeper to what would be expected by a response originating in the hand knob of the primary motor cortex. However, results from source localization should not be interpreted as the exact origin of the i-TEPs. First, the activity induced by the TMS could involve neurons over a few centimeters, while the position of the dipole is the result of the integration in one source of the activity of an entire cortical patch (Darvas et al. 2004). Second, more electrodes may be needed for a precise localization of i-TEPs of such high-frequency activity (Zelmann et al. 2014). Lastly, potential residual noise may result in localization error. At this latency, it is possible that residual decay artifact and/or small amplitude muscle artifact may still be present. Indeed, the muscle artifact is explained by a more anterior and deeper cortical source, as shown by our source localization results on individuals in which TMS induced a muscular response. Therefore, if a residual muscle artifact is present in the data, it may shift the source localization model toward this concurrent source. While the source localization models clearly distinguished between the generator of the i-TEPs and generators of muscle twitches, further studies therefore are needed to understand the exact cortical generator of i-TEPs.

Our investigation on the muscle artifact shows several important aspects related to immediate TMS-EEG activity. On one hand, it provides evidence that muscular activity does not contribute to the origin of i-TEPs and i-TRP as suggested from the results of the control experiment. The stimulation of a lateral site in correspondence with the temporal branch of the facial nerve close to FT7 started to evoke a clear muscular response on C3 even at very low TMS intensity, about 19% of the individual rMT (i.e., 10% of MSO). Increasing the TMS intensity to approximately 24% of rMT (i.e., 13% of MSO), the muscular response after nerve stimulation reached the amplitude of the response obtained when targeting M1 at 110% of rMT. At this extremely low intensity, it is unlikely that the underlying cortex was directly stimulated by TMS. If i-TEPs were contaminated by muscular artifacts, we would expect a similar pattern when delivering the stimulation in correspondence of a cranial nerve which elicits a clear muscular response, but likely no cortical response. While comparable in amplitude in the first 10 ms after the TMS pulse, the signal recorded from M1 and nerve stimulation showed marked differences both in the time domain as well as in the time-frequency pattern. Indeed, the three peaks characteristic of i-TEPs, clearly present after M1 stimulation, were absent after nerve stimulation. Consistently, the i-TRP showed an increased activity mostly at frequencies below 500 Hz. On the other hand, it is important to consider that the muscle artifact remains a major obstacle for recording immediate TMS-EEG responses. To give an idea, of the 28 participants collected in the original dataset, only half of them showed no muscle activity in TEPs recorded in the AP-PA condition, and even less (10/28) in the PA-AP condition. While online correction methods of the coil tilting have been suggested (Casarotto et al. 2022), the muscle is not an all-or-nothing type of response and optimized stimulations can still result in subtle muscle twitches that are difficult to spot. Considering that i-TEPs amplitude is around 30-60 microvolts, a very small muscle artifact may have the same amplitude. Therefore, this procedure may not be enough to completely remove its potential confounding effect.

To conclude, our findings converge in indicating immediate TMS-EEG responses as an excitability index of the stimulated motor cortex, ruling out the potential contribution of muscle artifacts. The characterization of a non-invasive measure of local excitability is crucial for advancing our understanding of various neuro-psychiatric conditions displaying cortical excitability alterations, and to develop personalized treatment strategies. Moreover, such evidence paves the way to future research non-invasively investigating the role of local activity on the signal propagation along the stimulated networks, possibly extending to other sensory and associative cortical areas.

