## Supplementary Materials for "Immediate TMS-EEG responses reveal motor cortex excitability"

Marta Bortoletto[1]#

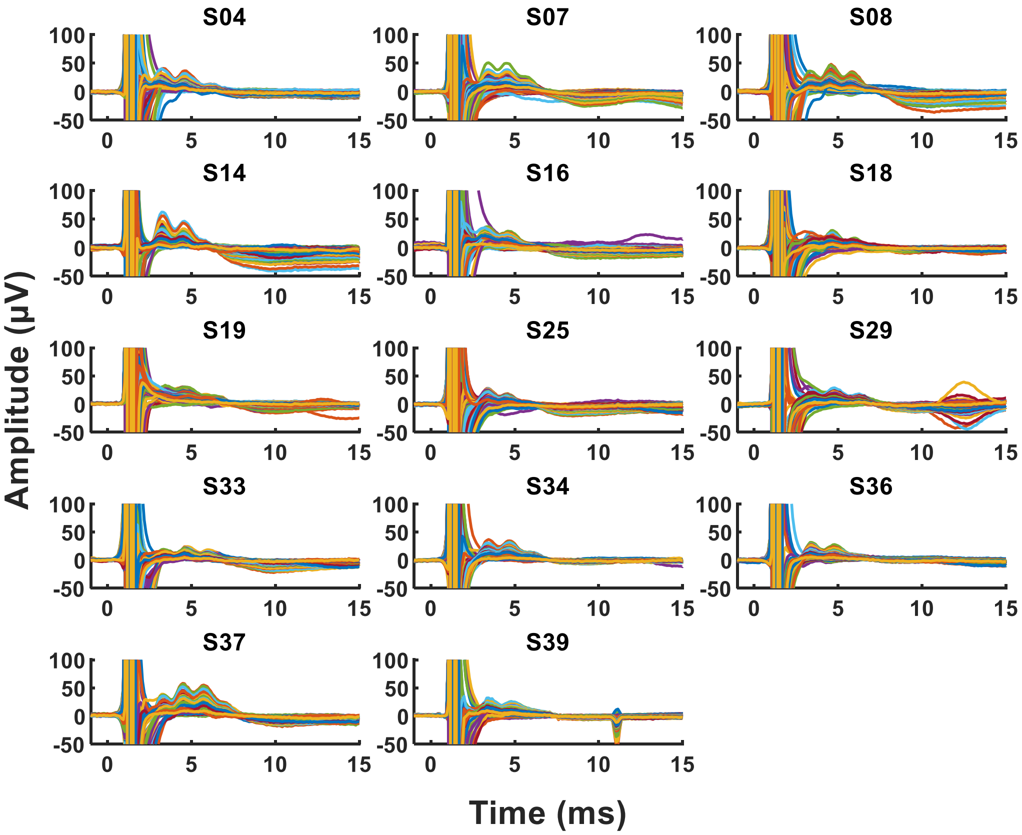


**Figure S1**. Individual TEPs average across trials for subjects of the NoMuscle group in the AP-PA condition.

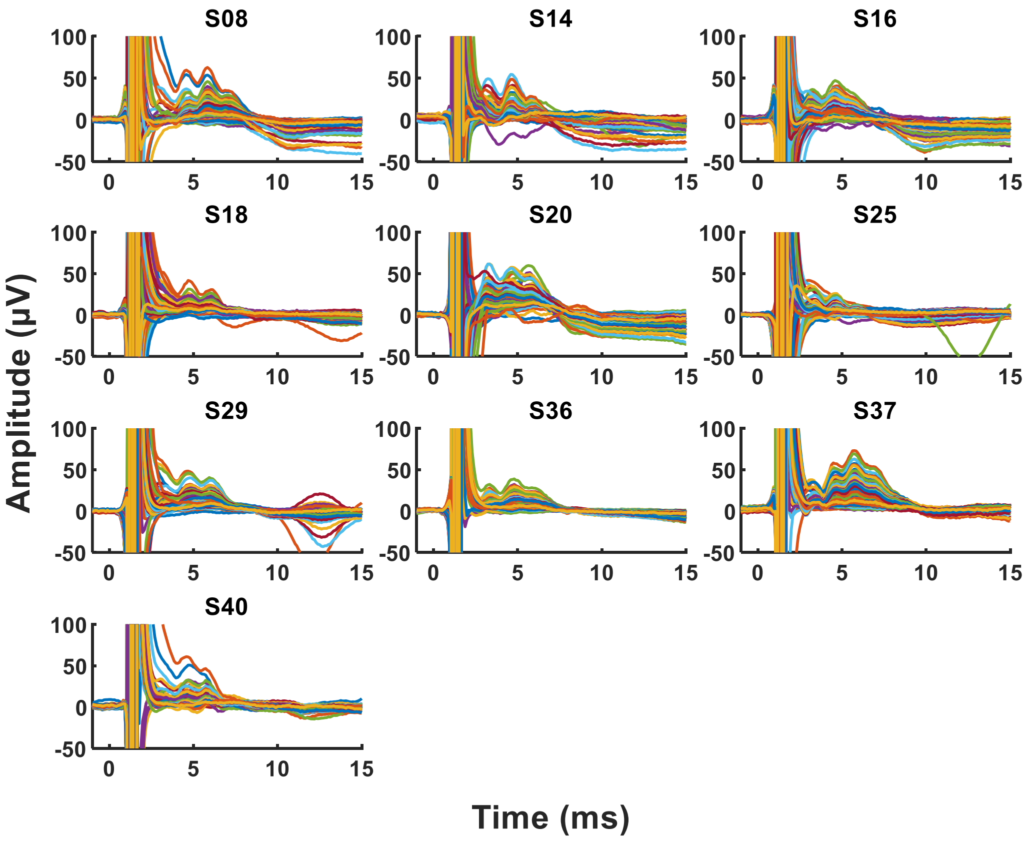


**Figure S2**. Individual TEPs average across trials for subjects of the NoMuscle group in the PA-AP condition.

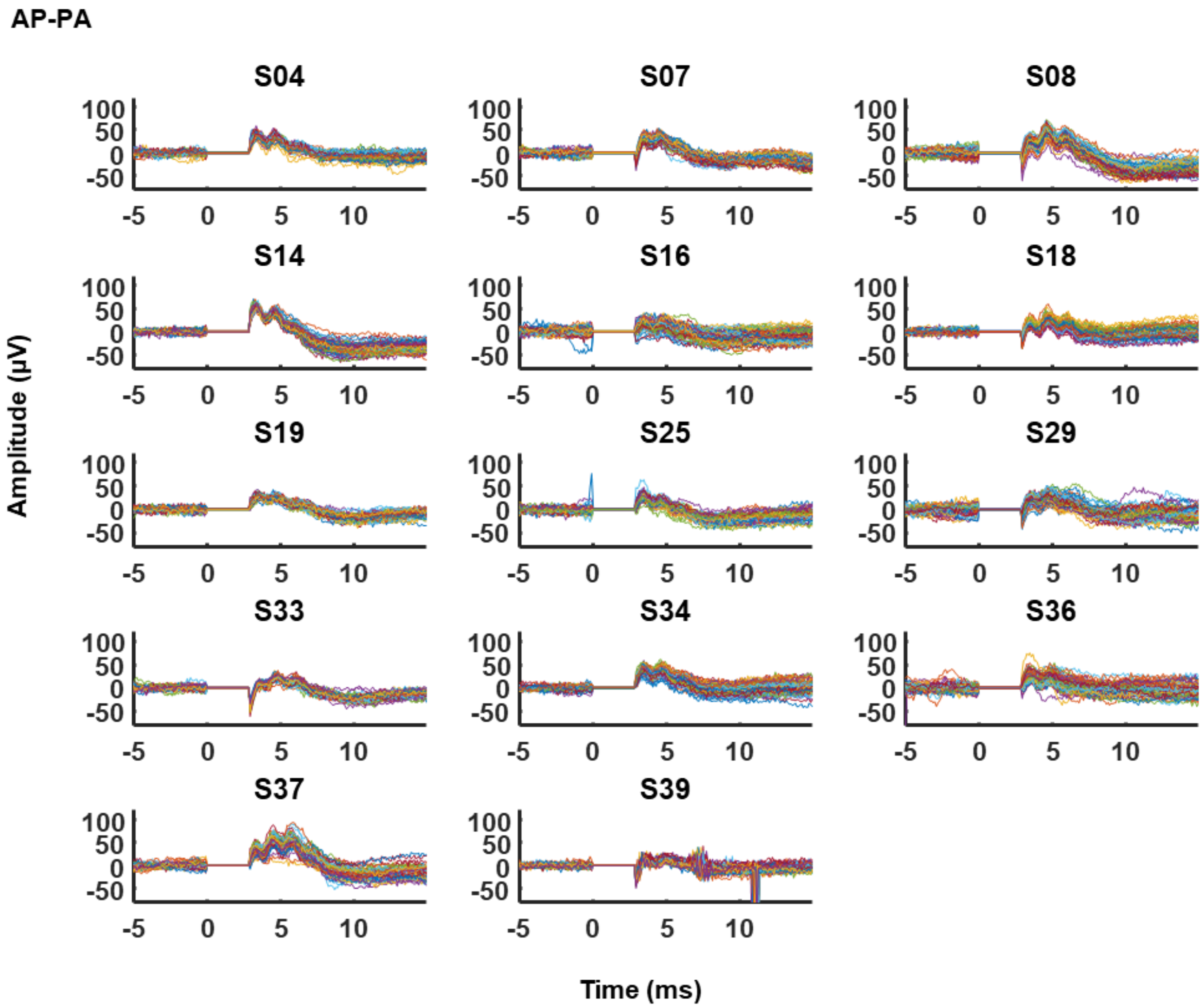


**Figure S3**. Single-trial iTEPs from individual subjects at channel C3 after artifact rejection in AP-PA. Each trace representing a trial. Baseline correction applied in the 10 ms before TMS pulse for display purposes.

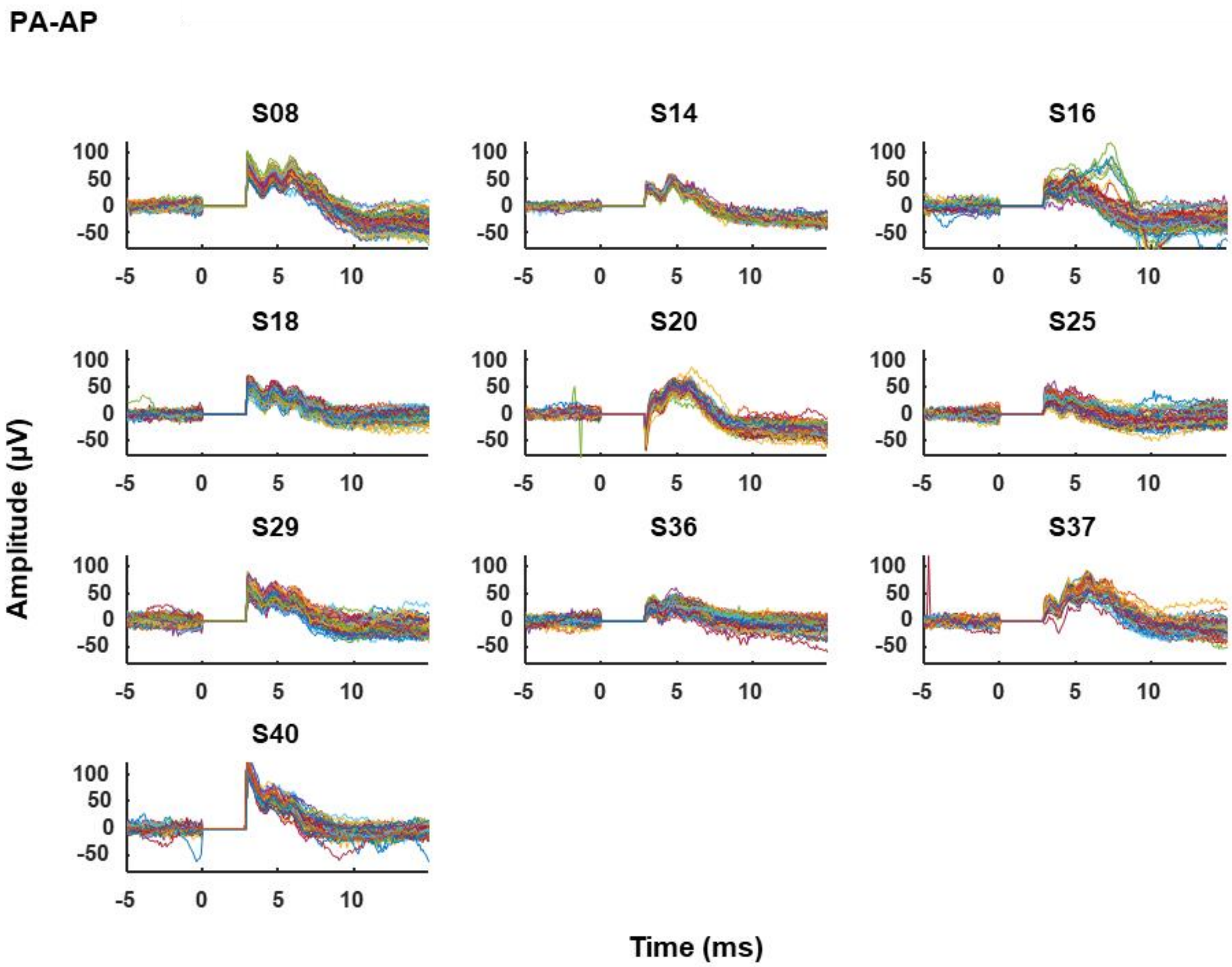


**Figure S4.** Single-trial iTEPs from individual subjects at channel C3 after artifact rejection in PA-AP. Each trace representing a trial. Baseline correction applied in the 10 ms before TMS pulse for display purposes.

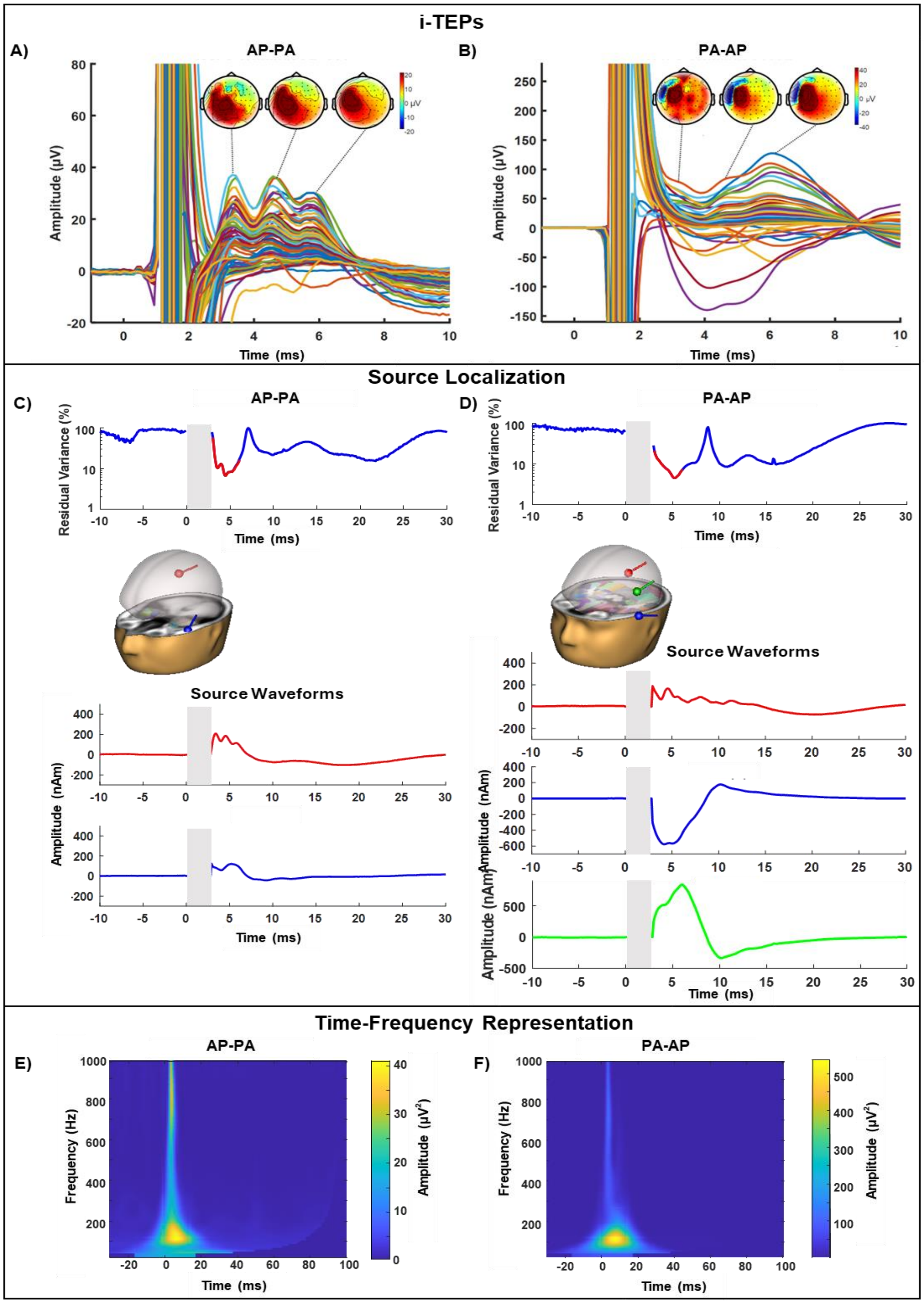


**Figure S5.** Grand-average across trials of the Muscle groups for AP-PA (panels A, C, E) and PA-AP (B, C, D) conditions. The top panels (A, B) depict the grand-average i-TEPs and the topography at the three peaks; the TMS artifact was interpolated in the analyses and is shown for display purposes. The middle panels (C, D) show the source localization of the dipoles that best explains the signal recorded over the scalp. The bottom panels (E,F) show the TFR at channel C3 (amplitude range in colorbar).

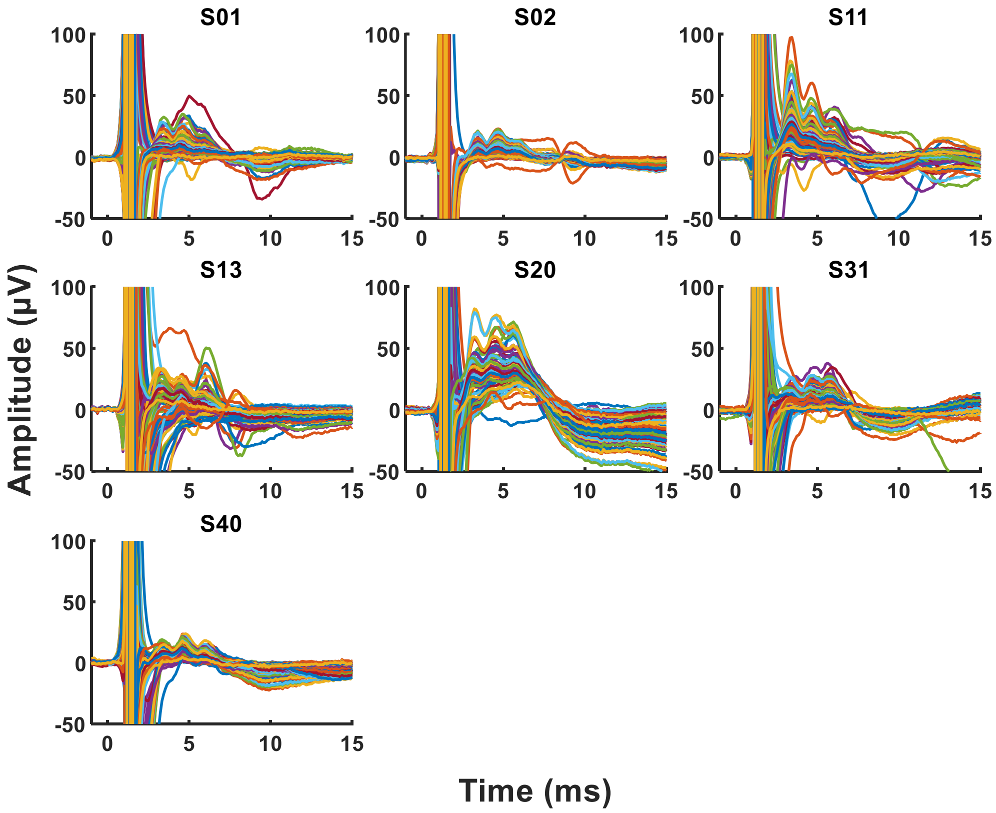


**Figure S6.** Individual TEPs average across trials for subjects of the Muscle group in the AP-PA condition.

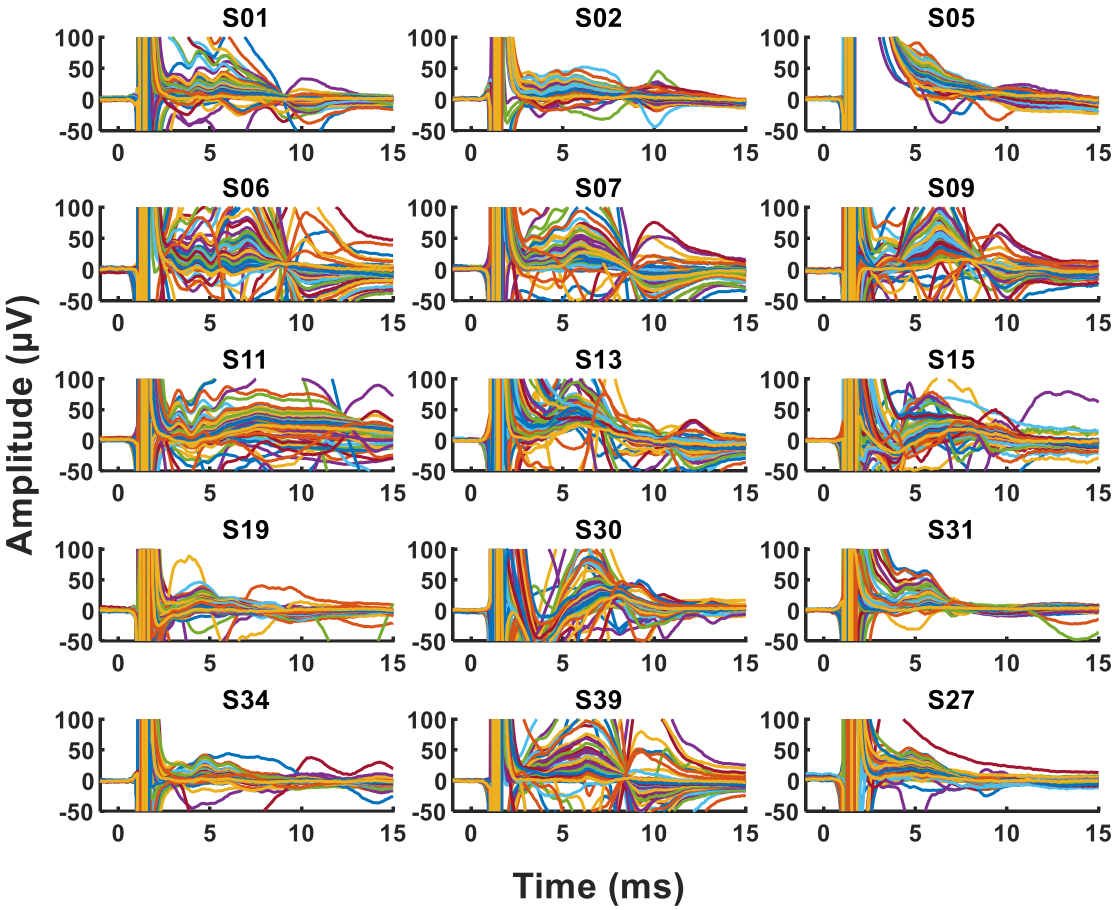


**Figure S7**. Individual TEPs average across trials for subjects of the Muscle group in the PA-AP condition.
